# Rearing condition affects behavioral response to cross-modal expectancy violation paradigm in zebra finches

**DOI:** 10.1101/2025.10.10.681716

**Authors:** Isabella Catalano, Sarah C. Woolley

## Abstract

Interacting in a multimodal world, including recognizing other individuals across multiple sensory modalities, is important for animals who rely on social relationships for survival. Social and sensory experiences shape how multisensory information is processed and integrated; however, we have less understanding of how multimodal recognition may be modulated by early sensory experience in a single modality. Zebra finches are gregarious songbirds that form lifelong pair-bonds with a single partner whom they recognize acoustically and visually. However, it is unknown to what extent multisensory signals might interact to enable recognition or whether this is affected by auditory exposure during development. In this paper, we tested females for responses to audio and visual stimuli from their mate or a stranger in a digital cross-modal expectancy violation paradigm. Using automated pose tracking, we determined that, like many species, zebra finches react differently when the stimuli are congruent versus incongruent. However, while birds reared in a colony setting and birds reared without exposure to adult song both detected audiovisual congruency, the degree of behaviors exhibited differed between the rearing conditions. Thus, multisensory integration appears to be important for females to identify their mates, but differences in developmental environment influence how recognition is behaviorally expressed.

## INTRODUCTION

Social living is critical to reproduction and survival across a diversity of species. Living in groups requires social animals to interact frequently, and many animals adjust their behavior depending on the identity of the individual they are interacting with. Thus, within a social group, interactions often depend on an ability to distinguish between individuals (1), whether to identify other group members or specific individuals (2, 3). Across species, the signals used for recognition span sensory modalities, including olfaction (4–7), audition (8–14), and vision (5, 15, 16). Additionally, many species integrate signals from multiple sensory modalities.

Multisensory integration is central to adaptive behavior and promotes individual recognition by taking advantage of the increased salience created by multisensory redundancy (17). The ability to form cross-modal representations of other individuals has been studied in a diversity of mammals, including lions, lemurs, and horses, as well as two species of birds (18–30). Much of the investigation of cross-modal individual recognition to date has focused on characterizing the presence or absence of the ability across species using a cross-modal recognition paradigm, which measures behavioral changes when a subject is presented with matching or non-matching stimuli; however, we know little about factors modulating multimodal recognition.

Social and sensory experience shape the way that audio and visual information are processed and integrated. During cross-modal integration, the degree of reliance on a unimodal sensory channel can depend on variation in sensory experience. For example, in humans, experience with a second language can affect cross-modal integration during speech perception. In particular, bilingual individuals show a greater McGurk effect, which is an indication of a greater reliance on visual information, compared to unilingual individuals (31). While these data highlight a potential role for sensory experience in multimodal sensory integration, the degree to which variation in developmental sensory experience in one domain might affect cross-modal learning and recognition is less well studied.

We investigated cross-modal recognition and the degree to which it is impacted by early sensory experience in zebra finches. Zebra finches are gregarious songbirds that live in large flocks and form lasting, pair-bonded dyads within those flocks (32). Hearing song during development shapes the way that female finches respond to song as adults. In particular, females reared without song exposure (“song-naïve”) do not show species-typical preferences or differential neural activation in response to courtship song compared to non-courtship song as adults (33, 34). In addition, while song-naïve females form pair-bonds with a male partner, their preferences for the mate’s song are not correlated with affiliative interactions in the same way as normally-reared birds (35). Adult, pair-bonded song-naïve females also lack the neural representation of the mate’s song that is seen in normally-reared birds (36). This last point raises the possibility that song-naïve females may differ in their integration of audio and visual information and their ability to cross-modally recognize their mate. Zebra finch females have previously been shown to identify their mates by song (37, 38) and by sight (16); however, whether they bind these two modalities to form a cross-modal representation of their mate is unknown. Here, we developed a novel, digitally-adapted version of a cross-modal expectancy violation paradigm to test both normally-reared and song-naïve female zebra finches for cross-modal recognition. Briefly, we presented pair-bonded female finches with a full-body video of their mate or an unfamiliar bird, followed by the screen going dark coupled with audio playback that was either congruent (audio and video from the same individual) or incongruent (audio and video from different individuals) with the individual in the video. Using machine-learning based pose-tracking software, we made detailed measurements of the bird’s movements and position to uncover a suite of behaviors that vary depending on congruency to demonstrate cross-modal recognition, and tested how these behaviors differ depending on a bird’s developmental sensory experience.

## METHODS

### Animals

Zebra finches (N = 35, n = 18 females, n = 17 males (1 male was paired twice), all >90 days post-hatch) had *ab libitum* access to seed, water, and grit and were kept on a 14:10 light:dark cycle. Egg and egg supplements were provided twice a week. Bird care and experimental procedures were approved by the McGill University Animal Care Committee and were performed in accordance with the Canadian Council on Animal Care guidelines.

Female birds were reared in one of two conditions. Normally-reared birds (n = 11) were reared with both parents and were exposed to the songs of other males in a colony setting, while song-naïve birds (n = 7) had their father removed 5-7dph, and were reared by their mother only in sound-attenuating boxes (TRA Acoustics, Cornwell, Ontario). Song-naïve birds remained in the nest until 60dph, at which point females were moved to a female-only colony setting.

### Experimental design

For all studies, female birds were housed with a male in a cage (10 in. x 10 in.) within a sound-attenuating chamber (“soundbox”). Within the same soundbox, a second male-female pair was housed in a different, adjacent cage. All pairs were given access to nesting material and an empty dish in which to nest. Birds were paired for at least two weeks, which previous research indicates is sufficient time to form a pair bond (35).

### Stimulus creation

Prior to mating, males were recorded singing both directed (female-oriented) and undirected song using Sound Analysis Pro (SAP; (39)) as previously described (34, 35, 38, 40). Briefly, during a song recording session, each male was presented with a stimulus female (a female bird otherwise not in the experiment) for 30 seconds to elicit directed song. After the male sang or after 30 seconds the female was removed and undirected song was collected. Interleaved recording of directed and undirected song continued until males had performed at least 10 songs in each condition. Songs were bandpass filtered (300-10 kHz), normalized by the maximum amplitude, and saved as a .wav file (44.1 kHz) using custom Matlab code (Mathworks, Natick, Massachusetts, USA). For each male, it was decided to use either directed or undirected song, and two song samples were utilized (3.35s average song length).

Video stimuli were collected using a GoPro Hero 7 (GoPro Inc., San Mateo, CA; narrow view, 120fps, 1080p). A white background was set up behind a T-shaped perch, and males were released into the space. The GoPro was set up at approximately eye-level from them, and recorded their entire body. Once the males were perched, video was recorded for approximately 1 minute.

Stimuli were assembled in VSDC Video Editor (Flash-Integro LLC., Toshkent, Uzbekistan). There were four stimulus conditions (Fig. 1): mate congruent (mate video, mate song), mate incongruent (mate video, unfamiliar song), unfamiliar congruent (unfamiliar video, unfamiliar song), and unfamiliar incongruent (unfamiliar video, mate song). Each video segment was played silently for 30s, at which point the screen went black and the corresponding song was played. Two song clips were played separated by 5s; on average, the total song exposure came out to 6.67s. Each condition (video + song) was separated by 5 minutes. In total, the stimulus was approximately 22 minutes long. Each bird was tested one time.

**Figure 1.**
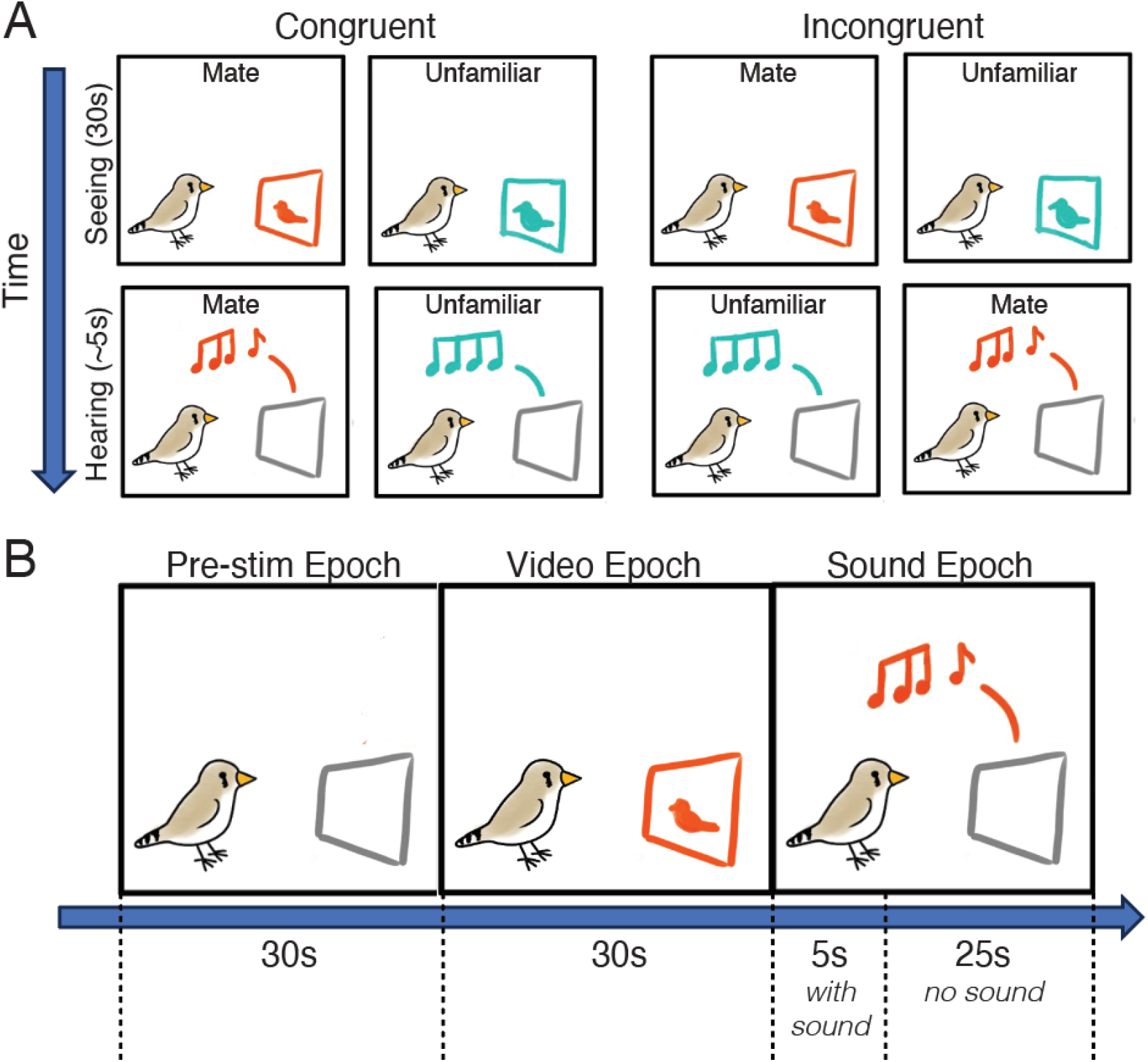
Experimental setup. A) Both congruent and incongruent stimulus conditions begin with a 30s video presentation of either the mate or the unfamiliar bird. After the video, the screen goes blank and subjects are played a song: in the congruent conditions, this song belongs to the individual they just saw, whereas in the incongruent conditions, it belongs to the opposite individual. B) Approximate timeline of the analyzed epochs. 30s of experimental video are taken before the video starts (pre-stimulus epoch), then 30s during the video playback (video epoch), and finally the 30s post-video, which contains approximately 5s of song towards the beginning and is thus called the sound epoch.

### Behavioral assay

Prior to testing, females were separated from their mate and housed alone in a soundbox overnight (for 18-24h) to acclimate to the testing chamber. A GoPro Hero 7 (superview, 60fps, 1080p) was set on top of the cage, looking down onto the bird, so that an overhead video could be recorded the next day. 15-30min before testing, water bottle and food dish were removed from the testing cage. A digital screen (ROG Strix XG17AHPE; ASUSTeK Computer Inc., Taipei, Taiwan) was set up inside the soundbox. The GoPro was set to record, and then the stimulus playback was initiated.

After the experiment, females were either reunited with their mate and moved to individual pair housing, or else divorced and returned to same-sex group cages. Exceptionally, one male was repaired and used as a mate for a second time.

### DeepLabCut analysis

Videos of the subjects’ behavior during stimulus presentation (henceforth ‘experimental video’) was divided into smaller segments using DaVinci Resolve (Blackmagic Design Pty Ltd., South Melbourne, Victoria, Australia). For each condition, experimental videos were divided into three epochs: 1) a ‘pre-stimulus’ epoch of 30s consisting of the 30s immediately preceding playback of the silent video; 2) a ‘visual’ stimulus epoch consisting of the 30s of playback of the silent stimulus video and 3) an ‘sound’ stimulus epoch which contained the song playback. Video recorded during the 5 minutes between stimulus presentation conditions was cut into ∼2min segments and used for training a DeepLabCut (DLC; (41)) model, detailed below.

For body part tracking, we used DeepLabCut version 2.3.10 (41–43). Specifically, we chose three body parts to label: the tip of the beak, the center of the bird’s head, and the center of the bird’s back. We labeled 20 frames taken from 21 training videos of 18 animals (then 95% of labelled frames was used for training). We used a dlcrnet_ms5-based neural network with default parameters for 3 training iterations. We validated with one shuffle, and found the test error was:

8.39 pixels, train error: 2.87 pixels (image size was 1080 by 1920 pixels). We then used a p-cutoff of 0.6 to condition the X,Y coordinates for future analysis. This network was then used to analyze experimental videos. After training, all 30s epoch videos (pre, video, sound) were analyzed for all stimulus combinations (3 epochs x 4 stimulus conditions gave us 12 videos per subject bird, 216 videos total) using DLC to acquire x and y pixel coordinates for every labelled body part (beak, head, back) for every frame of video.

### Data processing and statistical analysis

We quantified a number of aspects of behavior and movement during each epoch for all stimulus combinations. We manually counted the number of short vocalizations (‘calls’) produced in each experimental video epoch. We also processed DLC-generated x- and y-coordinate positions using custom-written Python 3.10.14 code. We focused on two aspects of movement and position: the overall position and movement of the bird and the direction and movement of the head. We used the x-coordinates of the bird’s back to calculate the means and standard deviation of the overall position relative to the screen and the linear variance (standard deviation squared) in proximity to the screen. To analyze the movement and direction of the head we took the x- and y-coordinates of the beak and head and used an arctan function to calculate the head angle relative to a vertical line in cartesian space (this vertical line could be mapped onto the left side of the experimental video). The calculated angle represents the direction of the bird’s head at any given moment (see Fig. 1C-D). From this, we calculated the mean, standard deviation, and circular variance (standard deviation squared) of the head angle using the Scipy package. For reporting, the location of the screen has been aligned to 0 degrees on the circle. We also determined the number of frames that the bird was facing in a particular direction. Specifically, we binned frames containing certain angles of absolute value into four categories: 0-20 degrees = binocular vision; 21-150 degrees = monocular vision; 56-65 degrees = ‘target’ monocular range; 151-180 = blind spot (where birds were unable to see the screen, and thus were looking away).

These bins were based on a previous study of zebra finch vision, which found 60 degrees to be a frequent fixation point for zebra finches (44). Any frames for which DLC did not provide a position were dropped, and the number of frames with values in each bin were taken as percentages of total frames with values.

There were four birds who did not look at the screen (defined as having a “target” looking range of 55-65° relative to the screen) in half or more of trials; these birds were dropped from results, as expectancy violation depends on the birds having seen the visual stimulus in the first place.

We used standard least squares mixed effects models with individual ID as a random variable and Tukey’s HSD for comparison tests unless otherwise noted. In some cases, birds did not have tracked frames during the first 2s of an epoch; these birds were dropped from that specific analysis (noted in Results). We tested whether there were differences in behavior between the rearing conditions or across the experimental epochs and whether differences across epochs varied by rearing condition. Within the video epoch we tested for differences in response by rearing condition and stimulus ID. Within the sound epoch we tested for effects of rearing, congruency, and audio stimulus ID. In all cases with more than one independent variable models were full factorial with all possible interactions. All statistical analyses were performed in JMP Statistical Processing Software (RRID:SCR_014242).

Finally, we tested the degree to which the category of the stimulus could be predicted by the bird’s behaviors we used a stepwise discriminant function analysis (DFA) with a leave-one-subject-out (LOSO) method of cross-validation to classify 1) video identity during the video epoch and 2) congruency during the sound epoch. Because the data set contains non-independent data points from the same subject (pseudoreplication; (45)) we tested for significance compared to a permuted DFA. For this, we performed within-subject permuted DFAs where the classification category was randomly assigned within each subject to generate null distributions. We report the accuracy of the DFA on the original data compared to the mean permuted DFA for

1000 permutations as a p-value where p < 0.05 indicates that the DFA classifies the data correctly significantly more often than the null model. The DFA was performed using custom-written Matlab code.

## RESULTS

We tested female zebra finches for cross-modal recognition by presenting them with a visual stimulus followed by a sound stimulus, where the video and sound were either congruent (from the same stimulus individual) or incongruent (from different stimulus individuals). Thus, there were four conditions: two congruent conditions (mate video followed by mate song; unfamiliar video followed by unfamiliar song) and two incongruent conditions (mate video followed by unfamiliar song, and vice versa; see Fig. 1A). Subjects were recorded overhead and recordings for each stimulus combination were split into three epochs: pre-stimulus epoch, consisting of the 30s before the video stimulus presentation; video epoch, consisting of the 30s during which the visual stimulus was presented; and sound epoch, the 30s immediately following the video epoch, of which the first few seconds was a song playback (Fig. 1B). We quantified a number of parameters of movement and attention including the direction that birds were looking relative to the screen, the number of vocalizations (“calls”), and the mean and variance of the overall position in the cage as well as the angle of the head for each epoch across all conditions.

### Overall movements within the arena vary depending on rearing condition

We first investigated the overall differences in behavior across the different epochs between the rearing conditions. We tracked the position of the bird relative to the screen and tested the degree to which there were changes in the amount that birds moved around the cage (positional variance) or in their overall proximity to the screen (mean position; see Methods). We found that normally-reared birds were, on average, closer to the screen than song-naïve birds across all epochs of testing (F(1,12) = 5.0552, p = 0.0441; Fig 2B). However, there was no significant difference in the variance, indicating that birds from both rearing conditions moved around in the arena similar amounts, but normally-reared birds did so in closer proximity to the screen.

**Figure 2.**
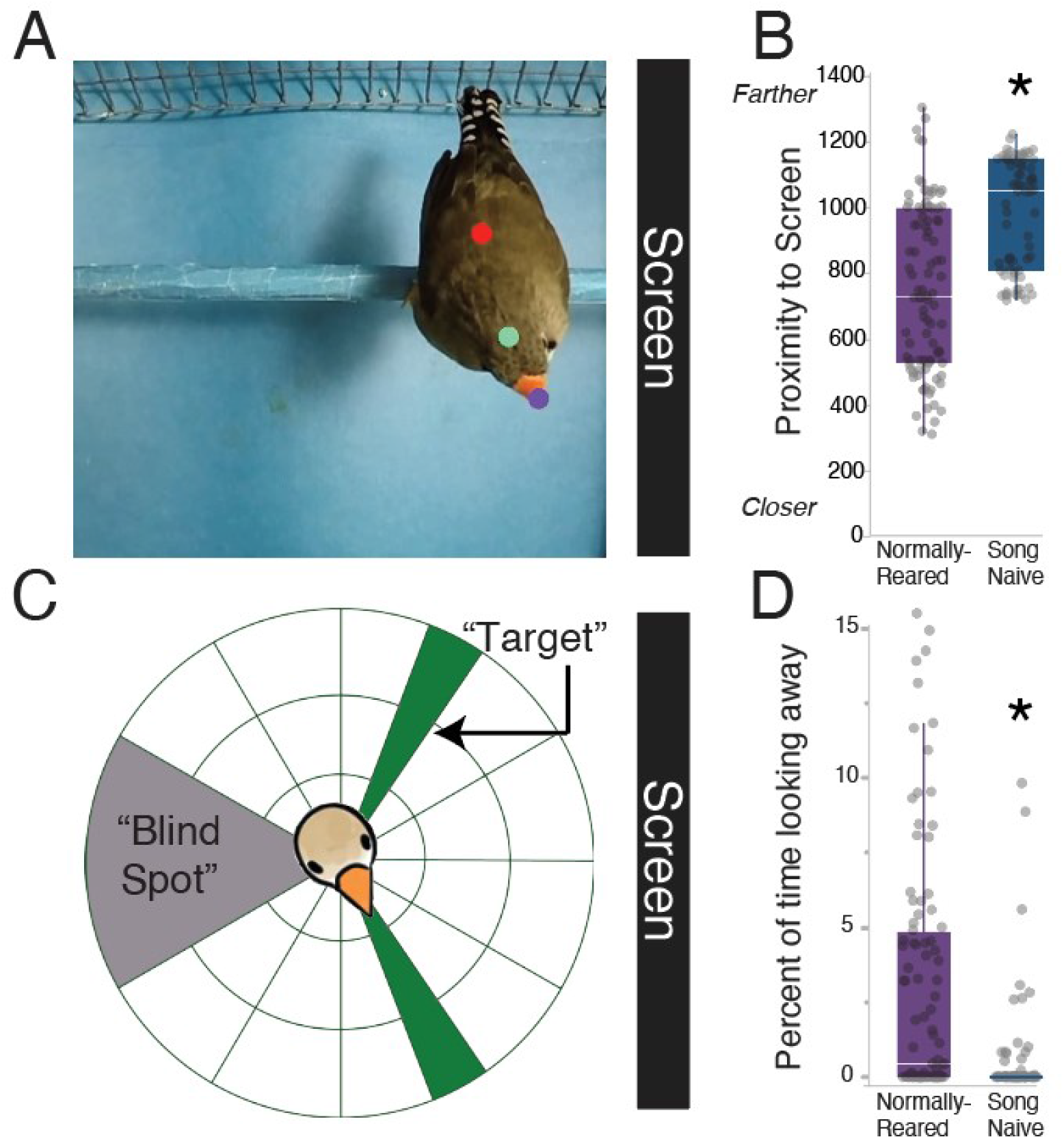
Overall movements differ between rearing conditions. A) Still image from an experimental video showing the subject bird relative to the screen. Colored dots are the three motion-tracking points used for pose analysis. B) Overall, across all epochs, normally-reared birds were significantly closer to the screen than song-naïve birds (p = 0.0441). C) Schematic illustrating head angles relative to the screen. The grey shaded area represents the birds’ “blind spot”, or an area in which they are facing away from the screen. The green shaded areas represent the “target” looking angle centered around 60°. The screen is always aligned to 0°. D) Normally-reared birds spent more time looking away from the screen than song-naïve birds (p = 0.0494). * indicates p<0.05

In addition, we calculated the direction of the bird’s head as an angle relative to the screen. Head angles of 55-65° indicate that the screen is within the bird’s target range of visual focus ((44); Fig. 2C, see Methods for more details). We also captured information on the angle and movement of the head by calculating the mean and circular variance of the head angle, with greater variance indicating more turning of the head. Normally-reared birds spent more time looking away from the screen (±151-180°) compared to song-naïve females (F(1, 12) = 4.7766, p = 0.0494; Fig. 2D), though there was no interaction or main effect of epoch.

### Zebra finches from both rearing conditions attend to visual and audio stimuli

We next verified that birds attended to the video playback. Previous work has shown that female finches produce short vocalizations (“calls”) during social interactions (9, 46) and in response to preferred songs (34, 47). While there was substantial variation across individuals in their call rates (0-86 calls, avg. 5.537 calls per video), we generally found that birds increased their calling during the social contexts (F(2, 150) = 9.4564, p = 0.0001, Fig. 3A). In particular, birds from both rearing conditions produced significantly more calls during both the video (p < 0.0001) and sound epochs (p = 0.0487) compared to the pre-stimulus epoch. There was no effect of rearing on call count (F(1,12) = 0.6551, p = 0.4340).

**Figure 3.**
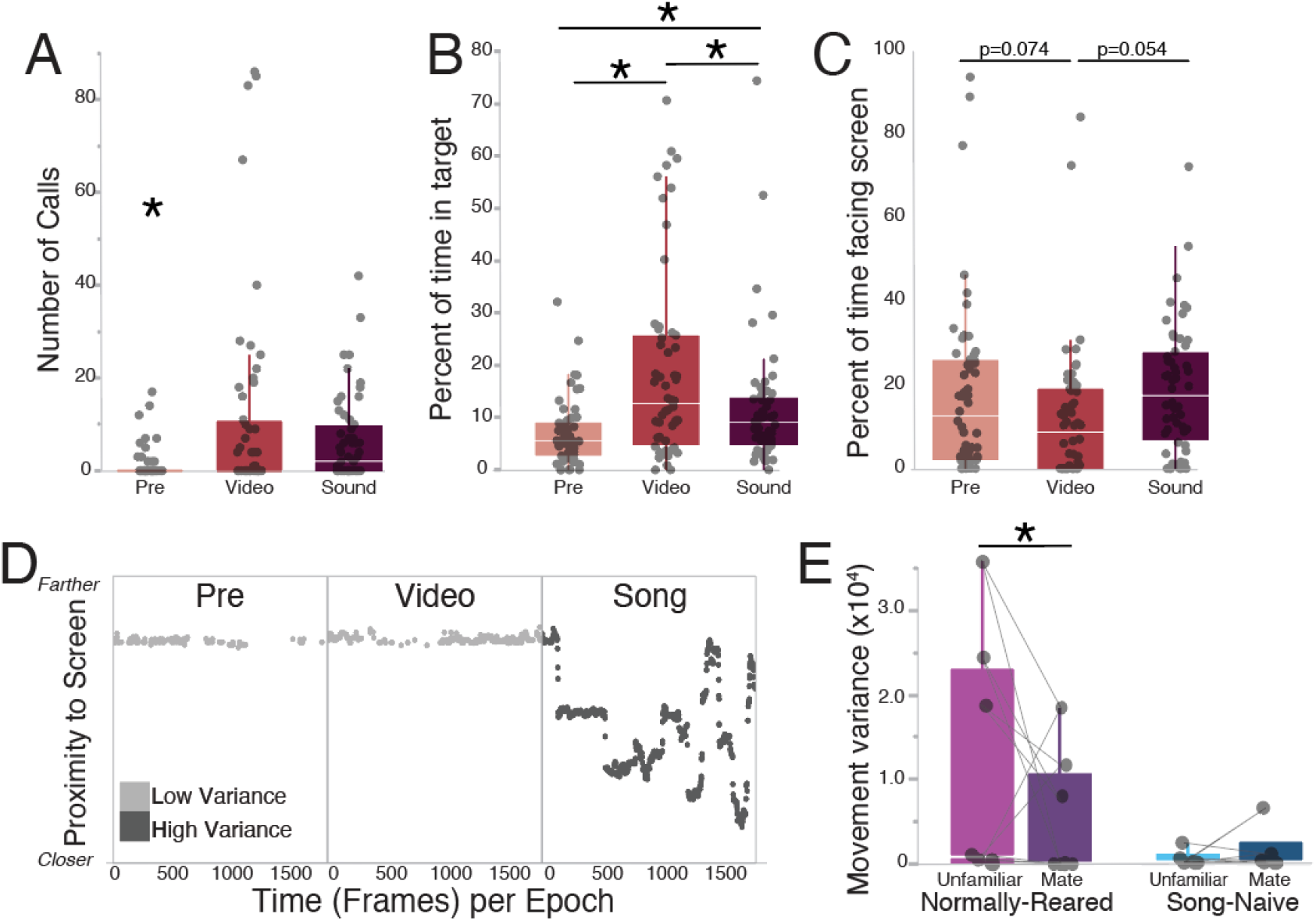
Behavioral indices vary by epoch. A) Birds significantly increased their calling during the video and sound epochs compared to pre-stimulus (p = 0.0001). B) Birds spent a greater percentage of time in the target looking range of 55-65° during the video epoch when there was a video on the screen than during the pre-stimulus or sound epochs (p < 0.0001). C) Conversely, birds spent less time in the binocular vision range of 0-20° during the video epoch compared to the pre-stimulus epoch (p = 0.0346). D) Data from an example bird plotting the position of the bird’s back relative to the screen (at the top). This bird demonstrated high positional variance during the song epoch only. E) There is an interaction of rearing and video stimulus for movement variance (p = 0.0222). Normally-reared birds moved more when shown a video of an unfamiliar bird instead of their mate, while song-naïve birds demonstrated no significant difference. * indicates p<0.05

We also investigated the degree to which birds look at the screen. Because the speaker is located in the screen, we tested whether birds adjusted their gaze direction during both video and audio playback. We found that all females, regardless of rearing, spent more time in the “target” looking range (±55-65° relative to the screen) during 30s of video presentation compared to when no video was present. Specifically, there was a main effect of epoch (F(2, 150) = 18.1719, p < 0.0001), with birds spending a greater percentage of time in the “target” range during the video epoch as compared to the pre-stimulus epoch (p < 0.0001) or the sound epoch (p = 0.0014; Fig. 3B) when no video was presented. Females also spent more time in the “target” range during the sound epoch compared to the pre-stimulus epoch (p = 0.0428). Conversely, there was a main effect of epoch on the time spent facing the screen (e.g. with the screen in the range of binocular vision 0-20°; F(2, 150) = 3.4405, p = 0.0346; Fig. 3C). Specifically, there was a trend for less time spent facing the screen during the video epoch compared to the pre-stimulus or sound epochs (p = 0.0736 and p = 0.0543, respectively). We did not find any significant differences between the epochs in the time spent looking away from the screen or in the overall amount of movement or mean position relative to the screen. Taken together, based on the changes in calling as well as the time in the target looking range, birds appear to attend to visual stimuli.

### Normally-reared birds differentially respond to a video stimulus depending on the identity of the stimulus

We next tested whether birds vary their behavior when the bird in the video playback is their mate compared to when the stimulus is an unfamiliar male. During the video epoch, the amount of movement within the arena varied depending on both the identity of the bird in the video as well as the rearing condition of the subject bird (F(1, 40) = 5.1233, p = 0.0291). Normally-reared females moved around the arena more (i.e. displayed greater movement variance) when they saw a video of an unfamiliar male than when presented with a video of their mate (p = 0.0222), while there were no differences within song-naïve females or between song-naïve and normally-reared females (p > 0.05 for all; Fig 3E). Within the video epoch, there was no difference in the mean head direction or the time in the “target” looking area, the time spent looking away, or the time spent facing the screen when birds were presented with their mate versus a stranger. There were also no significant differences in the amount of head movement (circular variance) either between rearing conditions or depending on the bird presented in the video.

### Zebra finches alter their movements depending on the congruency of audio and video stimuli

While a diversity of mammals show the ability to recognize other individuals using multiple sensory modalities, such multimodal recognition has not been widely tested in other taxa. We tested for cross-modal recognition and the degree to which it might be affected by developmental experience by analyzing behavior during song playback, the epoch when subjects were presented with a difference in congruency. We found significant effects of stimulus congruency as well as differences between the rearing conditions on the amount of body movement as well as the movement and direction of the head.

Within the sound epoch, birds from both rearing conditions were, on average, closer to the screen during incongruent playbacks compared to congruent playbacks (F(1, 40) = 7.5877, p = 0.0088; Fig. 4A). The closer proximity to the screen was especially prominent in birds that saw an unfamiliar video and then heard the mate compared to birds that saw and heard an unfamiliar male (p = 0.0142). During both congruent and incongruent conditions preceded by video presentations of the mate the positions of females relative to the screen were intermediate (p > 0.05 for all other pairwise comparisons). Thus, birds from both rearing conditions approach the screen specifically during the incongruent condition when they hear their mate after seeing an unfamiliar male.

**Figure 4.**
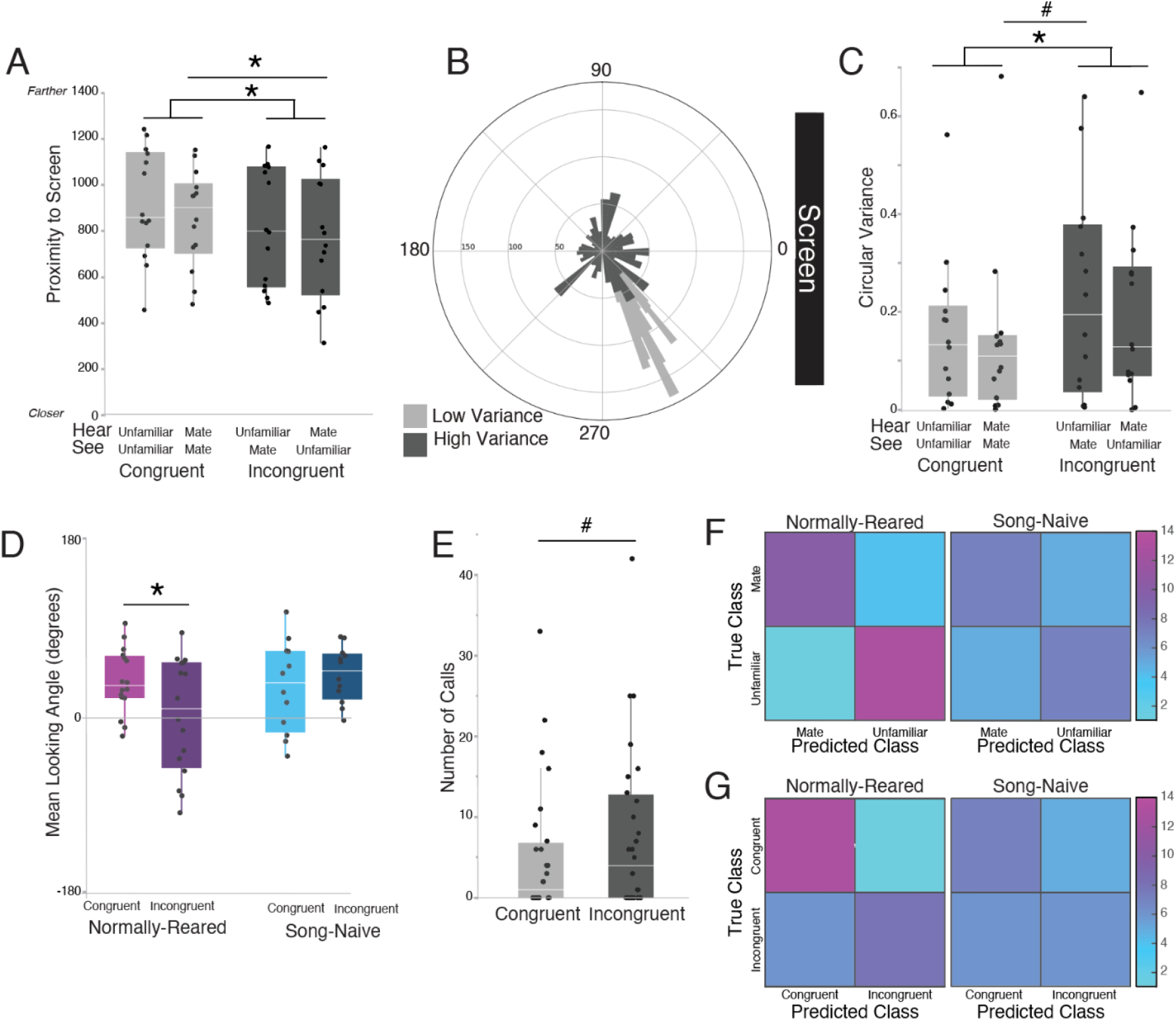
Body and head movements predict congruency. A) Birds from both rearing conditions were on average closer to the screen during incongruent conditions (dark grey; p = 0.0088). B) Schematic plotting the circular variance of two example birds. The head angle of the bird for each frame was sorted into ten-degree bins on a polar plot. One bird has low variance (light grey) and has a narrow range of bins; the other bird (dark grey) demonstrates high variance, with a more equal spread of angles around the circle. C) Circular variance during the first 5s of the sound epoch differed based on congruency, with incongruent conditions (dark grey) eliciting greater circular variance than congruent conditions (p = 0.0133). D) There was an interaction of rearing and congruency for the mean head direction (p = 0.0129), with normally-reared females showing significant changes in direction between congruency conditions but song-naïve females not doing so. The central line represents 0°, or the angle at which birds are facing the screen head-on. E) Females tended to call more during incongruent trials compared to congruent trials, though this was not significant (p = 0.0894). F) Confusion matrix from a discriminant function analysis (DFA) classifying the identity of the male showed in the video epoch based on the behavior of normally-reared (left) and song-naïve (right) females. G) Confusion matrix from a discriminant function analysis (DFA) classifying the congruency of the sound based on a subset of behaviors of normally-reared (left) and song-naïve (right) females. Normally-reared birds exhibit behaviors that can significantly predict the true class, but song-naïve females do not. # indicates p<0.10; * indicates p<0.05

In addition to approaching the screen, birds also altered their head movements (circular variance) and average head direction (circular mean) during incongruent compared to congruent stimuli (Fig 4B). Specifically, when songs were first played back, birds that were shown incongruent stimuli turned their heads more (had higher circular variance) than birds who were shown congruent stimuli (F(1, 40) = 6.7112, p = 0.0133; Fig. 4C; see Methods). There was a slight, but not significant, effect of stimulus order on the amount of head turning (F(3, 36) = 2.4399, p = 0.0802), with mate video followed by an unfamiliar song eliciting slightly greater circular variance than mate video followed by mate song (p = 0.0977).

The average direction that the head was facing (circular mean) was modulated by an interaction of rearing and congruency (F(1, 40) = 8.3105, p = 0.0063; Fig. 4D). Specifically, normally-reared females had mean head angles of 1 degree during incongruent condition, which is the angle when the head is facing the screen (located at degree 0), compared to -34 degrees during congruent conditions (p = 0.0129). Song-naïve females had a mean looking angle of -28 degrees for congruent and -41 degrees for incongruent conditions. Within song-naïve females and between song-naïve and normally-reared females there were no significant differences (p > 0.05 for all). Overall, coupled with variance in head movements, this suggests that birds are turning their heads from side to side frequently, such that the mean value is closer to the center.

The increase in head turning and changes to mean head position during incongruent trials were only present during the song playback. In particular, when we analyzed the circular variance over the remaining 25s of the sound epoch, there were no significant effects of the congruency of the sound playback to the previous video, or of the identity of the song on the circular variance (p > 0.05 for all).

Finally, while identity of the video did not have any significant effects on call count (p > 0.05 for all), there was a trend for females to call more during the sound epoch for incongruent condition trials compared to congruent condition trials (F(1,40) = 3.0300, p = 0.0894; Fig. 4E).

### Video identity and cross-modal recognition can be predicted based on behavior

We next investigated whether individual or combinations of behavioral parameters could be used to predict either the identity of the male in the video (mate vs. unfamiliar; Fig. 4F) or the congruency of the video and song stimuli (congruent vs. incongruent; Fig. 4G). We used a discriminant function analysis (see Methods) and, because of the variation in behaviors between the rearing conditions, we created separate models for normally-reared and song-naïve females. This allowed us to additionally investigate whether there were differences between the rearing conditions in which specific behaviors lead to greater predictive power.

We found that in normally-reared birds, the amount of head during the first 5 seconds of video playback correctly predicted the male in the video as the mate or an unfamiliar bird with 78.6% accuracy. Adding in the bird’s position relative to the screen brought the average accuracy to 82% with successful prediction of the mate (71%) and of the unfamiliar male (93%; p=0.0001 compared to the null model). In contrast, for song-naïve birds, the amount of movement around the arena provided the greatest predictive power with 58% mean accuracy. However, it did not accurately categorize the identity of the video significantly better than the null model (p=0.3590). Thus, as we saw with the individual parameters, normally-reared but not song-naïve birds altered their movements during video playback and these changes were predictive of the identity of the male in the video.

Both normally-reared and song-naïve females varied aspects of their behavior in response to a mismatch between audio and video playback. Interestingly, the behaviors that provided the greatest predictive power to determine whether a bird perceived a song to be congruent or incongruent differed between the rearing conditions. In normally-reared birds, a model with the average direction that the head faced during song playback (the circular mean), the percent of time spent looking in the target region, and the amount of head turning during song playback provided the most accurate predictions of whether a stimulus was congruent or incongruent (p<0.0001 over the null model). For song naïve birds, the position relative to the screen and the amount of movement in the arena resulted in the most accurate prediction of congruency. However, this model did not perform significantly better than the null model (p=0.4240).

## DISCUSSION

The ability to interact in a multimodal world is important for social animals who rely on complex social webs and relationships for their survival (1). Of particular importance for social animals is the ability to recognize another conspecific individual across modalities, a skill which has been experimentally demonstrated in a number of species, the majority of which have been mammals (18–30). While there is ample evidence for unimodal individual recognition in birds in both visual and auditory modalities (see (48) for review), only two cross-modal recognition studies, one involving crows (27) and the other penguins (20), have been performed to date. Here, we used a digitally-adapted version of an expectancy violation paradigm coupled with machine-learning-based pose tracking software and found that zebra finches show cross-modal recognition, further broadening the range of species that have demonstrated this skill.

Similar to previous work (16, 49, 50), we find that zebra finches attend to visual stimuli displayed on a screen. Both normally-reared and song-naïve females call more frequently and spend greater time with the screen in their target visual range when the video is playing compared to the epoch prior to playback. Additionally, we found that normally-reared birds responded differently depending on who was in the video, and these differences in behavior were sufficient to predict, with high accuracy, whether the video was of the mate versus an unfamiliar male. Thus, unlike previous work which required training of birds in order to test for visual recognition (16), we were able to leverage the natural changes in movements and behavior to uncover differential responses to video of the mate compared to an unfamiliar male.

In contrast to the responses of normally-reared birds, song-naïve females did not show any differences in behavioral response depending on the video identity. Moreover, there was not a model, based on the behavioral parameters that we measured, that predicted whether the video was of the mate or a stranger more accurately than chance. Although song-naïve females did not differentially respond to the mate versus the unfamiliar male in the video, this may not indicate a lack of recognition. Both song-naïve and normally-reared females discriminated between congruent and incongruent stimuli. In particular, when the video and sound did not match (were incongruent), females from both rearing conditions showed more head-turning behavior, consistent with literature on cross-modal individual recognition in other species. For example, changes in behavior between congruency conditions in other species include increased looking time in crows (27) and decreased sniffing of an olfactory cue in lemurs (28) during incongruent conditions. Discrimination of the congruency of audio and video requires the ability to recognize the mate during the video playback. Thus, although we do not see overt differences in their behavioral responses during video playback, song-naïve females must recognize their mate in the video. Whether song-naïve females do not vary their behavior in response to video of different males or, alternatively, that they show more subtle changes in behavior not captured by our analysis is unclear. Future experiments using higher-speed video to uncover differences in more subtle movements in response to the mate compared to an unfamiliar bird could test this possibility.

While congruency modulated the behavioral responses of both normally-reared and song-naïve females, there was variation in the strength of responses in the two rearing conditions. As with the video playback, the differences in behavior in response to congruent and incongruent stimuli were sufficient to predict congruency with high accuracy for normally-reared females but not in song-naïve females. Thus, our data highlight similar abilities in birds independent of acoustic rearing environment. Females from both rearing conditions recognize auditory and visual stimuli and are able to detect a mismatch in audiovisual representation. However, overall, we find differences in the complement or degree of behaviors seen in song-naïve females. For example, normally-reared birds were generally closer to the screen and had greater movement variance than song-naïve females. In addition, we were able to predict the identity of the bird in the video and the congruency based on behavior of normally-reared females but not song-naïve females.

That song-naïve females do not modulate the same behaviors or modulate them to the same degree depending on the identity of the male in the visual stimulus as normally-reared females builds on a previous study, which found that although song-naïve females formed species-typical pair bonds and preferences for their mate’s song, the strength of those preferences was correlated with different patterns of interactive behaviors compared to normally-reared females (35). Taken together, these results demonstrate the lasting effect of a diminished auditory experience during development on adult behavior. In particular, it seems that a lack of species-typical song and colony noise broadly impacts social behaviors. Zebra finches are a highly gregarious species where acoustic communication is key to social interactions. Future work further probing how the rearing environment during the first 60 days impacts social and motor behaviors and the neural circuits that could be underlying these differences will be key to understanding the degree to which early sensory experiences may not only shape sensory systems but have far-reaching effects on circuits critical for social behavior.

In the wild, zebra finches live in large flocks but form monogamous pair-bonds and mate for life, meaning they must repeatedly find the same individual among a large number of others. Thus, our demonstration that female zebra finches show cross-modal recognition of their mate fits with the natural history and social structure in this species. While our data indicate cross-modal representation of the mate, it is unclear whether zebra finches also form similar representations for other birds either in their family (e.g. their parents, siblings) or within a flock. Zebra finches have the capacity to discriminate between individual vocalizations from large numbers of individuals. For example, using operant training, birds can learn to discriminate up to 42 individuals based on their calls (14). However, the extent to which zebra finches can associate each of those calls with a visual representation of an individual is unknown. Further research expanding the question of individual recognition in zebra finches to encompass familiar individuals beyond the mate, such as neighbors or members of the same flock, would provide needed insight into the extent to which these birds use multimodal recognition in social interactions.

Furthermore, previous work in penguins has found distinct behavioral responses to a non-pair colony member, but not to the mate (20), with one possible explanation being that separation from the partner created a state of stress for the birds. Stress caused by mate separation has also been seen in zebra finches (51); our setup allows for 18-24h of habituation to being separated from the mate, but it is possible that our results nonetheless reflect a stressed state. Recent work suggests that zebra finch song serves to increase social cohesion in a flock (52), which necessitates individuals being able to distinguish neighbors from mate as well as strangers. A future study examining separation from the broader flock but not the mate specifically could serve to attenuate the stress response while further investigating multimodal recognition of neighbors or other familiar birds.

Their large vocal repertoires and extensive history in the world of neuroscientific research as established animal models make zebra finches an important species in the scientific world. In this paper, we have utilized a digital version of the expectancy violation paradigm, with the precision and control of stimuli that digitization enables, to show that zebra finches demonstrate the ability to recognize others cross-modally, and how environment during development can impact the behavior associated with this ability. The fact that zebra finches form dyadic pair-bonds while still living in a larger flock, combined with their extensive history of laboratory-based study, shows that zebra finches can be excellent model animals for studying individual recognition in the future.

## ACKNOWLEDGEMENTS

We would like to thank Kevin Zhang and Hannah Mosca for their assistance with data collection, and Kevin for his help with coding data organization. This work was supported by funding from the Natural Sciences and Engineering Research Council (NSERC RGPIN-2018-05267) and Canadian Institutes of Health Research (CIHR RN465106-469732) to SCW and fellowships from the Fonds de recherche du Québec–Nature et technologies (FRQNT 333009), Center for Research on Brain, Language, and Music (CRBLM), Healthy Brains Healthy Lives (HBHL), and the Max E. Binz Fellowship from McGill’s Faculty of Medicine to IC.

## DATA AVAILABILITY STATEMENT

Data and code will be made publicly available upon publication.

## AUTHOR CONTRIBUTIONS

IC and SCW designed the experiments; IC performed the experiments; IC and SCW analyzed the raw data; IC and SCW wrote and edited the manuscript.

